# Robust generation of erythroid and multilineage hematopoietic progenitors from human iPSCs using a scalable monolayer culture system

**DOI:** 10.1101/711267

**Authors:** Juan Pablo Ruiz, Guibin Chen, Juan Jesus Haro Mora, Keyvan Keyvanfar, Chengyu Liu, Jizhong Zou, Jeanette Beers, Hanan Bloomer, Husam Qanash, Naoya Uchida, John F. Tisdale, Manfred Boehm, Andre Larochelle

**Author notes:** Corresponding author: Andre Larochelle, M.D. Ph.D., National Heart, Lung and Blood Institute, National Institutes of Health, Bethesda, 9000 Rockville, Bethesda, MD 20892, USA. ☎ ✉ (301).

## Abstract

One of the most promising objectives of clinical hematology is to derive engraftable autologous hematopoietic stem cells (HSCs) from human induced pluripotent stem cells (iPSCs). Progress in translating iPSC technologies to the clinic relies on the availability of scalable differentiation methodologies. In this study, human iPSCs were differentiated for 21 days using STEMdiff™, a monolayer-based approach for hematopoietic differentiation of human iPSCs that requires no replating, co-culture or embryoid body formation. Both monolayer and suspension cells were functionally characterized throughout differentiation. In the supernatant fraction, an early transient population of primitive CD235a+ erythroid cells first emerged, followed by hematopoietic progenitors with multilineage differentiation activity in vitro but no long-term engraftment potential in vivo. In later stages of differentiation, a nearly exclusive production of definitive erythroid progenitors was observed. In the adherent monolayer, we identified a prevalent population of mesenchymal stromal cells and limited arterial vascular endothelium (VE), suggesting that the cellular constitution of the monolayer may be inadequate to support the generation of HSCs with durable repopulating potential. Quantitative modulation of WNT/β-catenin and activin/nodal/TGFβ signaling pathways with CHIR/SB molecules during differentiation enhanced formation of arterial VE, definitive multilineage and erythroid progenitors, but was insufficient to orchestrate the generation of engrafting HSCs. Overall, STEMdiff™ provides a clinically-relevant and readily adaptable platform for the generation of erythroid and multilineage hematopoietic progenitors from human pluripotent stem cells.

**Highlights:**

1. Robust, scalable and clinically-relevant monolayer-based culture system for hematopoietic differentiation of human iPSCs.
2. Successive emergence of primitive erythroid cells, definitive multilineage HSPCs and erythroid progenitors in the culture supernatant.
3. Abundant mesenchymal cells and limited arterial vascular endothelium in the culture monolayer.
4. CHIR/SB molecules increase arterial vascular endothelium formation, suppress primitive hematopoiesis and promote definitive multilineage and erythroid progenitors.

## 1. Introduction

Long-term repopulating hematopoietic stem cells (HSCs) have durable self-renewal properties and multilineage potential, and can thus serve as lifelong reservoir for all blood cells^1^. Hematopoietic stem and progenitor cell (HSPC) transplantation is the most established cellular replacement therapy for hematological and other conditions. However, the limited availability of HLA-matched donors and the associated severe acute and chronic complications significantly restrict allogeneic treatment options. In gene therapy applications, sufficient autologous HSPCs are not available for genetic modification in a wide variety of disorders, and methodologies for safe expansion of these rare cells are inexistent^2^. Induced pluripotent stem cell (iPSC)-based therapies are a promising alternative because of their potential to provide an unlimited source of autologous, patient-specific cells for transplantation. However, the stepwise addition of cytokines and morphogens used in most protocols to recapitulate the natural developmental process generally induces iPSCs to produce hematopoietic progenitors with limited engraftment and differentiation capability^3,4^. The self-renewal and multilineage capacity of iPSC-derived hematopoietic cells can be enhanced by forced ectopic expression of hematopoietic transcription factors (TFs)^5,6^. However, these protocols lack clinical relevance due to their low efficiency, poor scalability and the necessary use of integrating lentiviral vectors for constitutive expression of potentially oncogenic TFs.

In the developing mammalian embryo, blood is produced in consecutive waves^7-9^. The first wave of hematopoietic progenitors, dubbed “primitive”, occurs in the yolk sac blood islands^10^. This primitive program is highly restricted, resulting primarily in the emergence of large, nucleated CD235a+ erythroid cells expressing embryonic globins, and in the production of some macrophage and megakaryocyte progenitors. A second wave of hematopoietic potential, termed “definitive”, supplies lymphoid progenitor cells, as well as erythro-myeloid progenitors (EMPs) that produce red cells expressing predominantly adult-type γ- and β-globin molecules. The third wave is also considered “definitive” but is uniquely marked by the appearance of HSCs within the aorta-gonad-mesonephros (AGM) region of the developing embryo^11,12^. These cells are specified and bud off from hemogenic endothelium (HE), a distinct subset of vascular endothelium (VE) that directly changes fate into hematopoietic progenitors through a morphogenic process called endothelial-to-hematopoietic transition (EHT)^13-15^. Thus, HE with definitive hematopoietic potential must emerge for the ex vivo production of engraftable HSCs from human iPSCs.

Definitive multilineage HSPCs were shown to arise from HE lining arteries, but not veins, during development^16,17^. This observation evokes the possibility that arterial specification of HE is necessary to initiate the definitive HPSC program and exploitation of this concept may facilitate the ex vivo production of engraftable HSCs^18,19^. However, this hypothesis has been challenged and arterial specification of HE, while necessary, may not be sufficient for definitive HPSC formation^18^. In addition, distinct cellular signaling pathways regulate the specification of primitive and definitive hematopoiesis in human iPSC cultures. Repression of nodal/activin signaling and activation of the WNT/β-catenin pathway in mesodermal precursors are required to downregulate primitive and enrich for definitive hematopoietic programs^20-22^. Simultaneous modulation of these pathways at day 2 of culture has been shown to activate arterial genes and restore expression of several key hematopoietic regulators within the HOXA gene cluster that are commonly downregulated during human pluripotent stem cell differentiation^21,23^.

Most ex vivo methods for hematopoietic differentiation of iPSCs are based either on coculture with stromal cells or embryoid body (EB) formation^24^. Recently, an alternative simple, monolayer-based, chemically-defined, and scalable differentiation protocol requiring no replating, coculture or EB formation has become available (STEMdiff™ Hematopoietic Kit, STEMCELL Technologies, Inc.). During differentiation, a supportive adherent monolayer rapidly forms, followed by the emergence of suspension hematopoietic cells that can be harvested at regular intervals during culture for analysis. This approach reproducibly yields enriched populations of CD43+CD45+CD34+ hematopoietic progenitors with functional activity in the colony-forming unit (CFU) assay. The simplicity and efficiency of the protocol have enabled secondary applications, such as differentiation of microglia^25^ and NK cells^26^, validation of CD34 fluorescent reporter human iPSC lines^27^, mapping of human pluripotent stem cell differentiation pathways^28^, and investigation of a posttranscriptional regulatory circuitry for the maintenance and differentiation of pluripotent stem cells and HSPCs^29^. In this study, we systematically examined the phenotype and function of supernatant and monolayer cells throughout differentiation, and provide evidence that this system can be adapted to move closer to a clinically-relevant protocol for the generation of definitive erythroid and multipotent hematopoietic progenitor cells.

## 2. Methods

### 2.1. Generation and culture iPSCs

Peripheral blood CD34+CD38^-^ cells mobilized from healthy donors were reprogrammed into iPSCs (MCND-TENS2 line) using the integration-free CytoTune 2.0 Sendai virus reprogramming kit (A16517, Thermo Fisher Scientific) following the method previously reported^30,31^. Pluripotency was confirmed by teratoma formation assay, G-banding karyotype and flow cytometry for TRA-1-60 and NANOG markers as previously described^32^. iPSCs were maintained on 6-well tissue culture plates thinly coated with Matrigel® (Corning, 354230) in Essential 8™ (E8) Medium (A1517001, ThermoFisher). Culture medium was changed daily and iPSCs were split every three to four days. Briefly, medium was aspirated and wells were sequentially washed with phosphate buffered saline (PBS) or 0.5 mM EDTA in PBS (PBS/EDTA). Cells were then dissociated in PBS/EDTA 0.5 mM for 2-3 minutes. PBS/EDTA was aspirated and replaced with 1 mL E8 medium containing 1.25 µM ROCK inhibitor (Y0503, Sigma). Cells were pipetted 3-4 times using a P1000 pipette to dissociate into small to medium sized clusters, and split onto new 6-well plates at various dilutions.

### 2.2. Hematopoietic differentiation of human iPSCs

iPSCs were differentiated for 21 days using the STEMdiff™ Hematopoietic Kit (05310, STEMCELL Technologies, Inc.). Briefly, one day (D) before differentiation (D-1), iPSCs were split as described above, and cluster concentrations were calculated. A total of 20-35 clusters were transferred per well into a Matrigel-coated 12-well plate and cultured overnight. On D0 of differentiation, medium A was added to promote mesodermal differentiation and a half medium A change was done on D2. In select experiments, WNT/β-catenin agonist CHIR99021 (SML1046, Sigma) and activin/nodal/TGFβ antagonist SB431542 (S4317, Sigma) were added on D2 at a final concentration of 3 µM for 24-36 hours. On D3 of differentiation, supernatant was removed and hematopoietic differentiation medium B was added, followed by half-medium change on day 5, 7, 10, 12, 14, 17, and 19. On days when medium was changed, supernatant and monolayer cells were harvested for analysis. To completely detach the monolayer and clusters of cells from the plastic surface, Accutase™ (07920, STEMCELL Technologies, Inc.) was added to each well for 10 minutes at 37°C and cells were vigorously pipetted up and down to ensure complete recovery. Cells are then filtered through a 40 µM cell strainer, counted, and prepared for further analysis.

### 2.3. Flow Cytometry and Fluorescence-Activated Cell Sorting (FACS)

Cells were stained with antibodies (Table S1), following manufacturer’s instructions and analyzed on an LSRII Fortessa analyzer (Becton Dickinson). For gene expression, CFU assays, MSC differentiation, and mouse transplantation studies, cell populations were sorted on a FACS Aria (Becton Dickinson) with a 100 µm nozzle.

### 2.4. Gene Expression by real-time qPCR

RNA was extracted and purified using directions of the RNeasy Micro Kit (74004, Qiagen). To assess HOXA gene expression, real-time qPCR was done using TaqMan RNA-to-Ct 1-Step Kit (4392653, Applied Biosystems) on a BioRad C1000 touch system and all samples were multiplexed to include an internal *GAPDH* control. For evaluation of globin expression, real-time qPCR was performed using the Quantstudio 6 system (ThermoFisher), with an α-globin reference gene and a plasmid containing alpha-theta-epsilon-gamma-beta sequences in a 1:1:1:1:1 ratio for normalization of expression, as previously described^33^. Primers and probes used for real-time qPCR are listed in Table S2. Error bars were calculated using SEM for ΔΔCT values to calculate the upper and lower boundaries of the 95% confidence interval, which were then log transformed in the same way as the mean (2^-ΔΔCT^). Two-way unpaired Student’s t-tests were done on untransformed ΔCT values.

### 2.5. CFU Assays

Human CFU assays were performed as per manufacturer’s instructions (04445, STEMCELL Technologies, Inc.). Briefly, 9,000 and 4,500 total supernatant cells for control and CHIR/SB samples, respectively, were suspended in 300 uL of DMEM/2% FBS, which was then added to the methocult tube and vortexed. Following 5 minutes to allow for bubble dissipation, 1.1 mL of the medium was plated onto 35 × 10 mm style tissue-culture dishes (353001, Corning) for 1,500 to 3,000 cells per plate. Colonies were scored 17-20, and 14 days following plating for control and CHIR/SB samples, respectively.

### 2.6. NSG mouse transplantation

6-to 12-week-old female NSG mice (Jackson Laboratory, stock #05557) were sublethally irradiated (270 cGy) 24 hours before tail-vein injection of 3 × 10^4^ to 3×10^5^ CD43+CD45^+/-^ supernatant cells harvested and sorted at day 10 and 12 of iPSC differentiation. Bone marrow (BM) was collected 16 weeks post-transplantation and stained with human CD45-PE (clone HI30, BD Pharmingen). Animals were housed and handled in accordance with the guidelines set by the Committee on Care and Use of Laboratory Animals of the Institute of Laboratory Animal Resources, National Research Council (DHHS publication No. NIH 85-23), and the protocol was approved by the Animal Care and Use Committee of the National Heart, Lung, and Blood Institute.

### 2.7. Mesenchymal stromal cell (MSC) differentiation

To confirm MSC identify, CD43^-^CD45^-^ monolayer cells were sorted at various days of culture and differentiated per manufacturer’s instructions into adipocytes (A1007001, Thermo Fisher Scientific), chondrocytes (A1007101, Thermo Fisher Scientific), and osteocytes (A1007201, Thermo Fisher Scientific). For comparison, control MSCs derived from the bone marrow of a healthy individual were differentiated following the same procedures.

### 2.8. Erythroid differentiation

For determination of globin chain composition, supernatant and monolayer cells were collected at day 7 and 17 of culture, and differentiated toward the erythroid lineage as previously described^33,34^. Briefly, cells were cultured on irradiated OP9 cells (American Type Culture Collection (ATCC)) in Medium B with the addition of 5 U/mL erythropoietin (EPO; Amgen) and 5 ng/mL interleukin-3 (IL3; R&D Systems). Two days later, the floating cells were transferred into freshly irradiated OP9 feeder plates using an erythroid expansion media based on Iscove’s Modified Dulbecco’s Medium (IMDM; Sigma Aldrich) supplemented with 10 ng/mL stem cell factor (SCF; R&D Systems), 1 ng/mL IL3, 2 U/mL EPO, 1 µM estradiol (Pfizer), 1 µM dexamethasone (VETone, Boise), and 20% Knockout Serum Replacement (KSR, Thermo Fisher Scientific). Five days later, the medium was changed to a maturation erythroid medium containing IMDM, 2% bovine serum albumin (BSA; Roche), 0.56 mg/mL transferrin (Sigma Aldrich), 2 mM L-glutamine (Thermo Fisher Scientific), 2 U/mL EPO, 10 ng/mL insulin (Lilly), and 20% KSR, and the cells were cultured for another 8 to 10 days.

### 2.9. Reverse phase-high performance liquid chromatography (RP-HPLC)

The globin protein content of differentiated erythroid cells was evaluated by RP-HPLC as previously described^33,35^. Briefly, erythroid cells were harvested and washed 3 times with PBS followed by a lysis step using HPLC grade water. The lysates were mixed with 10% of 100 mM Tris (2-carboxyethyl) phosphine (TCEP, Thermo Fisher Scientific) and incubated for 5 minutes. Then, the lysates plus TCEP were mixed 1:1 with a solution containing 0.1% trifluoroacetic acid (TFA) and 32% acetonitrile (Honeywell Burdick & Jackson). The mixtures were centrifuged at 16,000g for 5 minutes and the supernatant was injected in an Agilent 1100 HPLC (Agilent Technologies) equipped with a reverse phase column, Aeris 3.6 µm Widepore C4 200Å (25034.6 mm, Phenomenex) with two solvents: solvent A, 0.12% TFA in water and solvent B, 0.08% TFA in acetonitrile using a flow of 0.7 mL per minute for 50 minutes.

### 2.10. Statistical Analysis

Results were analyzed with GraphPad Prism Software, using unpaired Student t-tests. Results are displayed as mean ± SEM and * signifies p<0.05, ** p<0.01, *** p<0.001, and **** p<0.0001.

## 3. Results

### 3.1. Robust production of an adherent monolayer and supernatant hematopoietic cells during differentiation of human iPSCs

Human iPSCs reprogrammed from CD34+ cells of a healthy volunteer (Fig. S1, A to C) were subjected to hematopoietic differentiation for 21 days using the STEMdiff™ monolayer approach. In select experiments, we also explored the effects of adding the WNT/β-catenin agonist CHIR99021 (CHIR) and the activin/nodal/TGFβ antagonist SB431542 (SB), from differentiation day 2 to 3 (Fig. 1A). Under culture conditions that favored mesodermal specification (medium A, day 0 to 3), an adherent monolayer rapidly formed. With the subsequent addition of hematopoietic cytokines (medium B, day 3 to 21), hematopoietic clusters emerged from the monolayer before their eventual release in the supernatant fraction (Fig. 1B).

**Figure 1.**
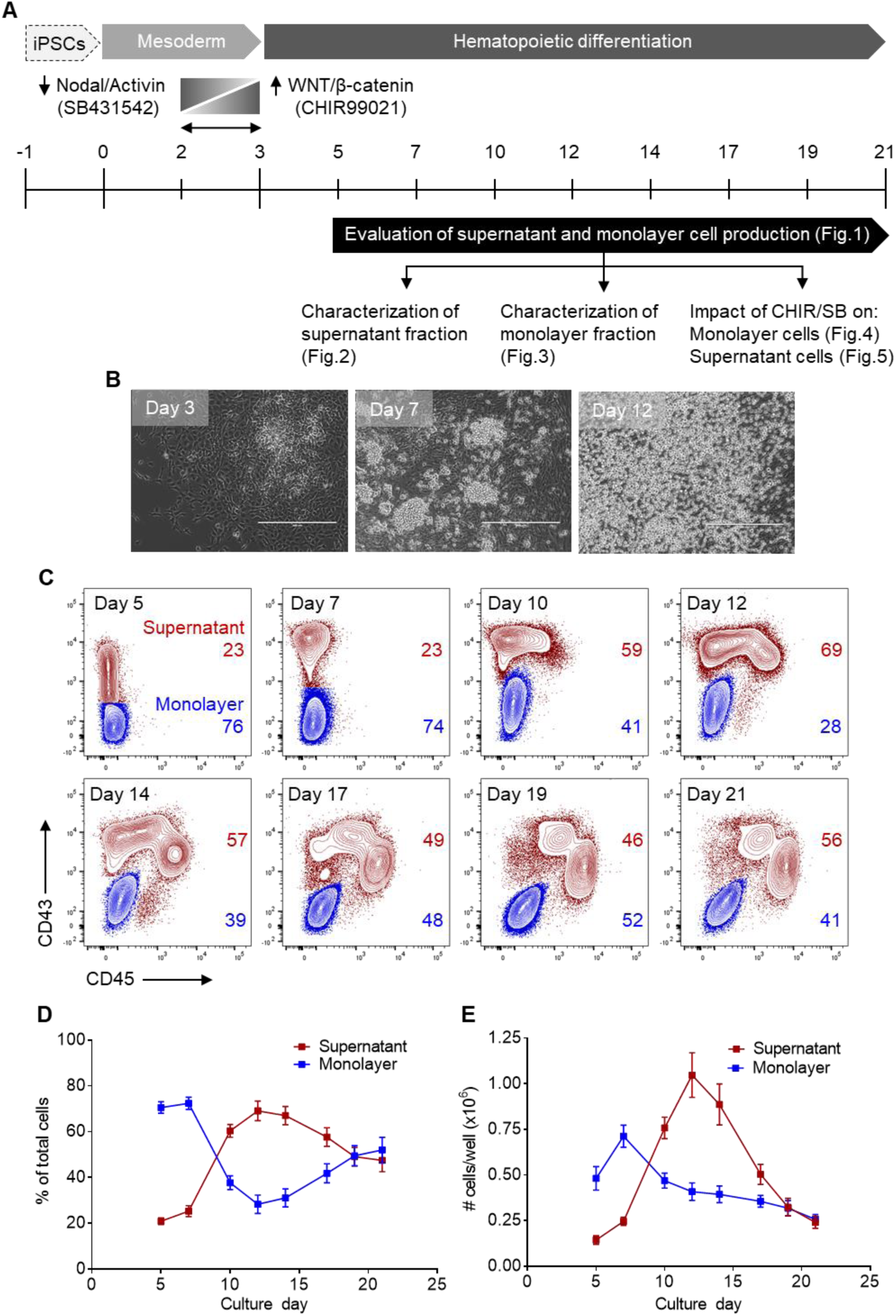
Robust production of an adherent monolayer and supernatant hematopoietic cells during differentiation of human iPSCs. (**A**) Schematic of the experimental design for hematopoietic differentiation of human iPSCs, including mesodermal (D0 to D3) and hematopoietic (D3 to D21) differentiation steps. Long vertical bars represent full medium changes or initiation/end of culture, and short vertical bars indicate half medium changes. Monolayer and supernatant cells were harvested for analyses at the indicated intervals from D5 to D21 of culture. In select experiments, WNT/β-catenin agonist CHIR99021 (CHIR) and activin/nodal/TGFβ antagonist SB431542 (SB) were added on D2 for 24-36 hours. (**B**) Representative phase-contrast microscopy images taken at various days of differentiation: D3-Adherent monolayer formation with the emergence of mesodermal colonies; D7-Hematopoietic clusters arising from the adherent monolayer; D12-Suspension hematopoietic cells. Scale bar = 400 µm. (**C**) Representative flow cytometry overlay plots of CD43 and CD45 expression, depicting CD43^-^CD45^-^ monolayer cells (blue) and CD43+CD45^+/-^ supernatant cells (red) at various days of differentiation. (**D**) Percentages of supernatant and monolayer cells at various days of differentiation (n=6). (**E**) Absolute numbers of supernatant and monolayer cells arising from 20-35 iPSC clusters (one well of a 12-well plate) at various days of differentiation (n=6). In panels (D) and (E), results are displayed as mean ± SEM.

To systematically characterize the supernatant and monolayer cells arising from this system, both populations were collected at regular intervals between day 5 and 21 of differentiation (Fig. 1A). To circumvent the technical challenges and inaccuracies associated with the physical separation of the supernatant and adherent fractions, both cell populations were harvested together at each time point and distinguished from each other by flow cytometry. Supernatant cells were characterized by varying expression of CD43, the earliest marker of human hematopoietic commitment, and by gradual acquisition of the pan-hematopoietic marker CD45 (CD43+CD45^+/-^). In contrast, cells of the adherent monolayer formed a distinct population expressing neither markers (CD43^-^CD45^-^) (Fig. 1C).

In the first phase of differentiation (day 5 to 9), the monolayer comprised most cells in culture with a modest relative contribution from the supernatant fraction (Fig. 1D, E). Between day 10 and 14, we observed an increase in percentages (Fig. 1D) and absolute numbers (Fig. 1E) of hematopoietic cells, and a concomitant decrease in the proportion of adherent cells likely resulting from an EHT process in culture. Supernatant hematopoietic cells were detected at a peak frequency of 69.1 ± 4.3% with an average yield of 1.1 ± 0.1×10^6^ cells per 20-35 iPSC clusters (one well of a 12-well plate) at day 12 of culture. In the third stage of culture (day 15 to 21), both hematopoietic and monolayer fractions contributed equally (Fig. 1D), but at a reduced total output (Fig. 1E). These data collectively show robust differentiation of human iPSCs into a supportive adherent monolayer from which hematopoietic clusters and suspension cells develop for up to 21 days in culture.

### 3.2. Successive emergence of hematopoietic populations in the culture supernatant during differentiation of human iPSCs

To determine whether hematopoietic differentiation of human iPSCs using the STEMdiff™ monolayer system can provide a suitable ex vivo model to study the emergence of hematopoiesis, we characterized the hematopoietic CD43+CD45^+/-^ supernatant fraction throughout differentiation by flow cytometry and functional assays.

We observed the formation of 3 successive populations of cells, as defined by expression of the CD34 hematopoietic progenitor cell marker. The first hematopoietic cells arising in the supernatant were predominantly CD34^-^CD45^-^ with a distinct CD235a+ erythroid phenotype. These cells peaked at day 7 and rapidly disappeared, evoking the transient nature of primitive erythroid cells that form during wave one of hematopoiesis in the developing embryo (Fig. 2A, B). The second population of suspension cells was enriched in CD34^hi^CD45^lo^ multipotent HSPCs. The maximum production of these cells was observed at day 12 of differentiation, comprising 17.8 ± 0.6% of the supernatant cells and producing 1.7 ± 0.2 × 10^5^ HSPCs per well (Fig. 2A, B). In the third phase of hematopoietic production, HSPCs gradually lost CD34 and upregulated expression of surface CD45, consistent with their progressive differentiation in extended cultures. By day 17 of culture, CD34^lo/-^CD45+ cells accounted for 81.9 ± 1.9% of the supernatant fraction, with an average yield of 3.7 ± 0.2 × 10^5^ cells per well, and this contribution persisted through day 21 (Fig. 2A, B).

**Figure 2.**
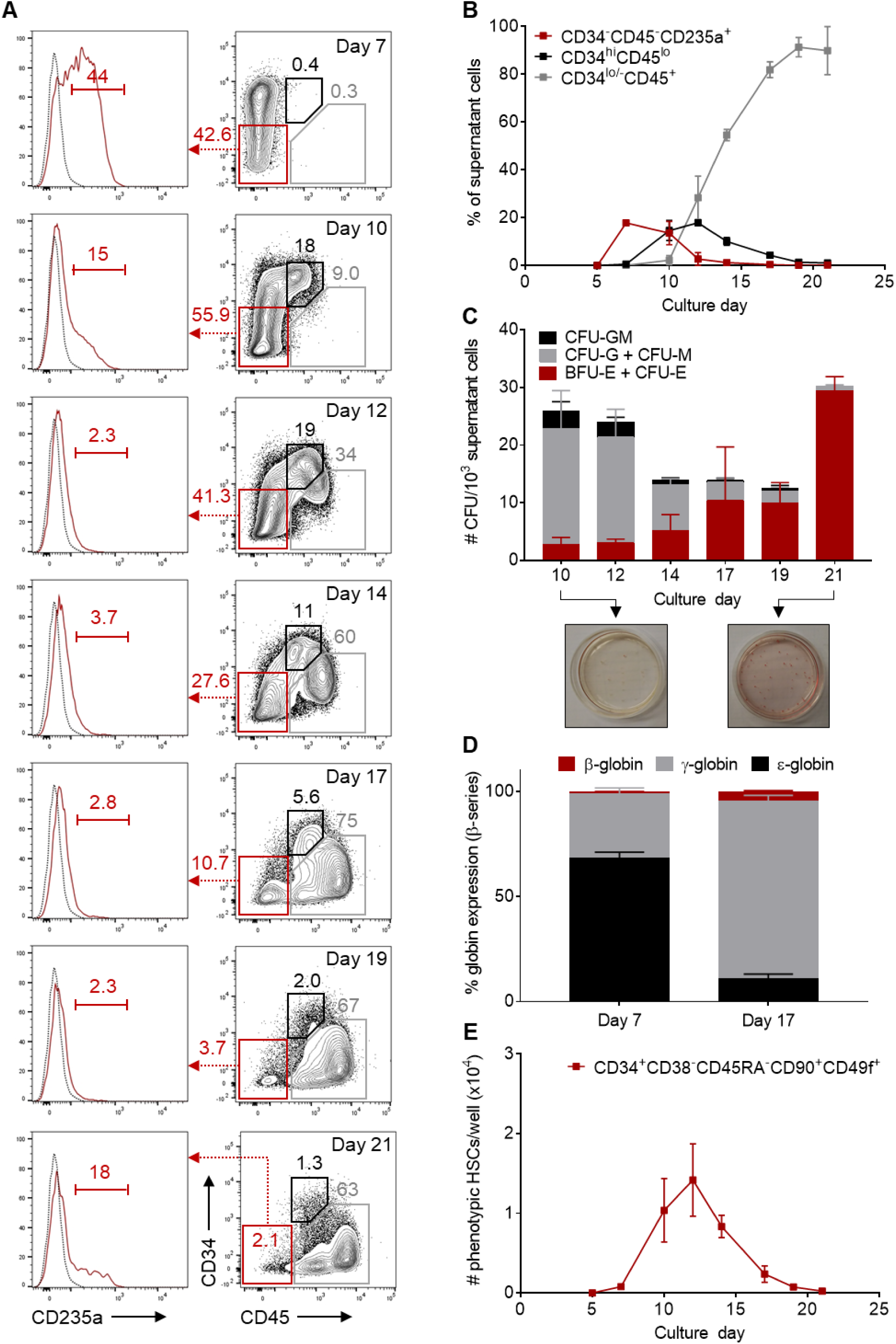
Successive emergence of primitive and definitive hematopoietic populations in the culture supernatant during differentiation of human iPSCs. (**A**) Representative flow cytometry plots of CD34, CD45 and CD235a expression in gated CD43+CD45^+/-^ supernatant cells at various days of differentiation. Gates shown include: CD34^-^CD45^-^CD235a+ primitive erythroid cells (red), CD34^hi^CD45^lo^ multilineage HSPCs (black), and CD34^lo/-^CD45+ hematopoietic progenitors (gray). (**B**) Percentage of hematopoietic populations with immunophenotypes classified above within the supernatant fraction at various days of differentiation (n=3). (**C**) CFU counts per 1000 sorted supernatant cells at various days of differentiation (n=6). Insets at the bottom are representative CFU plates demonstrating multilineage HSPCs at D10 and almost exclusive production of erythroid progenitors at D21 of culture. (**D**) Percentage of β-globin series expression in erythroid cells differentiated from D7 and D17 supernatant cells (n=3). (**E**) Absolute numbers of CD34+CD38^-^CD45RA^-^CD90+CD49f+ cells arising from 20-35 iPSC clusters (one well of a 12-well plate) at various days of differentiation (n=3). In panels (B) to (E), results are displayed as mean ± SEM.

We next sorted CD43+CD45^+/-^ supernatant cells to assess their multilineage differentiation potential in clonogenic progenitor assays. Colony-forming progenitors were detected throughout differentiation at frequencies ranging from 12.5 ± 4.8 to 30.3 ± 2.2 progenitors per 10^3^ supernatant cells (Fig. 2C). At culture days when CD34^hi^CD45^lo^ HSPCs were found to be most abundant by flow cytometry (day 10 to 14), both myeloid (CFU-G, CFU-M and CFU-GM) and erythroid (BFU-E and CFU-E) colony growth was observed. A progressive rise in BFU-E/CFU-E colonies was subsequently detected during culture, with a nearly exclusive erythroid contribution by day 21 (Fig. 2C). To evaluate the developmental stage of erythroid progenitors, we quantified globin chains by RP-HPLC in RBCs differentiated from day 7 and day 17 supernatant cells (Fig. 2D). At day 7, embryonic (ε) globins prevailed, comprising 68.4 ± 2.6% of total β-globin series expression. In contrast, red cells derived from day 17 suspension cells predominantly expressed adult-type γ- and β-globins, whereas ε-globins contributed much less compared with day 7 (11.1 ± 2.1% of β-globin series expression). These results were consistent with primitive (first wave) and definitive (second wave) hematopoiesis at day 7 and 17 of culture, respectively.

To determine whether the supernatant fractions at different stages of differentiation might contain hematopoietic cells with long-term repopulating potential, we first searched for cells with a CD34+CD38^-^CD45RA^-^CD90+CD49f+ phenotype which enables the highest reported purity of human HSCs^36^. Phenotypically defined HSCs (Fig. S2A) were detected in the culture supernatants primarily between day 10 and 14, with a maximum yield of 1.4 ± 0.5 × 10^4^ cells per well at day 12 of differentiation (Fig. 2E). However, these cells did not result in efficient, long-term engraftment after transplantation in immuno-deficient (NSG) mice, consistent with the previously reported dissociation between stem cell repopulating function and cell surface phenotype in cultured cells (Fig. S2B, left panel)^37^.

From these data, we infer that hematopoietic differentiation of human iPSCs with STEMdiff™ enables the sequential development of hematopoietic cells with features of primitive wave one-hematopoiesis (peak at day 7), definitive multilineage HSPCs with potent colony formation activity in vitro but limited engraftment potential in vivo (peak at day 12), and definitive erythroid-committed progenitors expressing adult-type globin chains (peak at day 17 to 21).

### 3.3. Abundant mesenchymal cells and limited arterial VE in the culture monolayer during human iPSC differentiation

To identify possible causes for the lack of durable repopulating potential of iPSC-derived HSPCs in this system, we sought to identify the cellular constituents of the CD43^-^CD45^-^ adherent monolayer from which hematopoietic cells develop during differentiation. Because definitive HSPCs emerge from HE in close association with the VE^10,11^, we first queried whether cells of the monolayer expressed VE-defining markers, including VE-cadherin (CD144) and high levels of surface CD34 (CD144+CD34^hi^). Non-hematopoietic cells expressing neither markers (CD144^-^ CD34^-^) were recently shown to characterize a population of mesenchymal cells^38^.

We found that most cells comprising the adherent population throughout differentiation lacked CD144 and CD34 expression, representing up to 92.5 ± 2.5% of the monolayer at day 21. In contrast, CD144+CD34^hi^ VE accounted for only a limited proportion of the monolayer, with a maximal contribution of 7.8 ± 2.0% at day 5 that steadily declined during culture (Fig. 3A, B).

**Figure 3.**
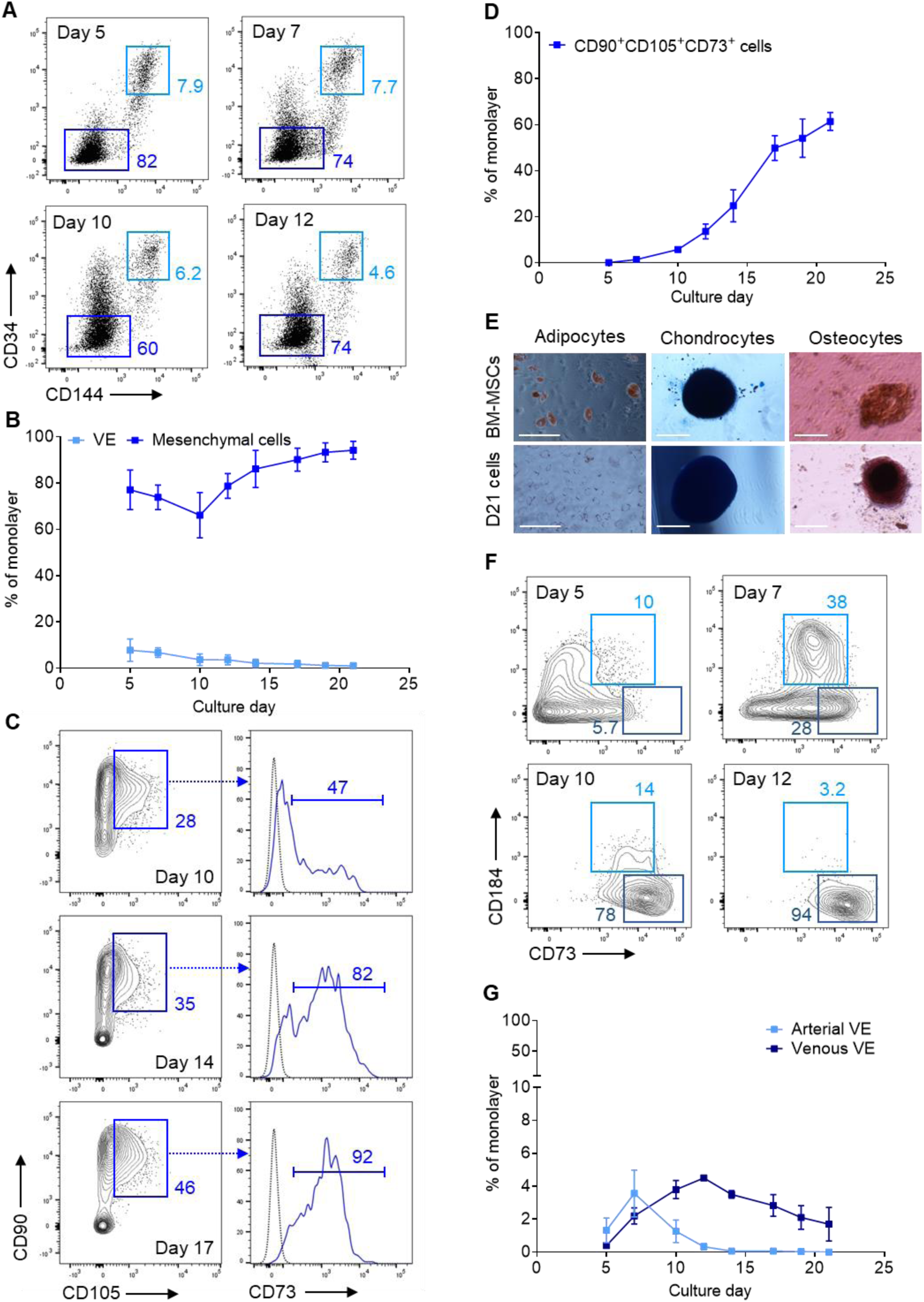
Abundant mesenchymal cells and limited arterial VE in the culture monolayer during human iPSC differentiation. (**A**) Representative flow cytometry plots of CD34 and CD144 (VE-cadherin) expression in gated CD43^-^CD45^-^ monolayer cells at various days of differentiation. Gates shown include: CD34^hi^CD144+ vascular endothelium (VE, light blue), and CD34^-^CD144^-^ mesenchymal cells (dark blue). (**B**) Percentage of cells with a VE or mesenchymal immunophenotype within the monolayer fraction at various days of differentiation (n=3). (**C**) Representative flow cytometry plots of CD90, CD105 and CD73 expression in gated CD34^-^ CD144^-^ mesenchymal cells at various days of differentiation. (**D**) Percentage of cells with a CD90+CD105+CD73+ mesenchymal stromal cell (MSC) immunophenotype within the CD43^-^ CD45^-^ monolayer fraction at various days of differentiation (n=3). (**E**) Representative images depicting adipocyte (Oil Red O stain), chondrocytes (Alcian Blue stain), and osteocytes (Alizarin Red stain) after culture of control bone marrow-derived MSCs (top panels) or D21 sorted CD43^-^ CD45^-^ cells (bottom panels) under conditions promoting differentiation of each cell type (n=2). Scale bar = 100 µm (adipocytes panels) or 200 µm (chondrocytes and osteocytes panels). (**F**) Representative flow cytometry plots of CD184 and CD73 expression in gated CD34^hi^CD144+ VE cells. Gates shown include: CD73^mid^CD184+ arterial VE (light blue), and CD73^hi^CD184^-^ venous VE (dark blue). (**G**) Percentage of cells with an arterial or venous VE immunophenotype within the monolayer fraction at various days of differentiation (n=3). In panels (B), (D), and (G), results are displayed as mean ± SEM.

To further characterize the CD144^-^CD34^-^ fraction, we assayed for expression of the surface markers CD90, CD73, and CD105, commonly used to identify mesenchymal stromal cells (MSCs), a major component of the adult BM niche. We observed a progressive increase in the percentage of cells with an MSC phenotype during differentiation (Fig. 3C, D), contributing up to 61.5 ± 3.9% of the monolayer at day 21. Because MSCs characteristically differentiate to adipocytes, chondrocytes and osteocytes in vitro, we tested the trilineage differentiation potential of day 21 monolayer cells. In agreement with the observed MSC phenotype, we detected adipocytes, chondrocytes and osteocytes, although differentiation to adipocytes was more limited compared to control MSCs derived from primary BM specimens (Fig. 3E).

Because HE and HSPCs are associated with arteries, not veins, in the developing embryo^19,39,40^, we next investigated whether arterial endothelium could be identified within the CD144+CD34^hi^ VE population. Based on distinct expression of the surface markers CD184 and CD73, arterial and venous VE progenitors were previously reported to segregate to the CD73^mid^CD184+ and CD73^hi^CD184^-^ fractions, respectively^41^. We observed that CD73^hi^CD184^-^ venous VE accounted for most of the CD144+CD34^hi^ endothelial fraction of the monolayer throughout differentiation. In contrast, low percentages of CD73^mid^CD184+ arterial VE were detected only in the first phase of differentiation, with a maximal contribution of 3.6 ± 1.4% of the monolayer at day 7 of culture. These cells subsequently declined and became undetectable after day 12 of differentiation (Fig. 3F, G). Thus, the limited arterial VE and the distinct abundance of mesenchymal cells within the adherent monolayer fraction could provide a sensible explanation for the lack of engraftment potential of iPSC-derived HSPCs in this system.

### 3.4. CHIR/SB molecules increase arterial VE in the adherent monolayer during the early phase of human iPSC differentiation

Because simultaneous modulation of nodal/activin and WNT/β-catenin pathways in mesodermal precursors was previously shown to activate arterial genes, restore expression of the hematopoietic HOXA gene cluster and promote definitive hematopoiesis^21,23^, we examined the effect of supplementing the culture medium with the WNT/β-catenin agonist CHIR99021 (CHIR) and nodal/activin inhibitor SB431542 (SB) at day 2 of STEMdiff™ iPSC differentiation for a period of 30-36 hours (Fig. 1A).

Addition of CHIR/SB led to a global increase in the percentages (Fig. 4A) and numbers (Fig. S3A) of CD43^-^CD45^-^ adherent monolayer cells on most days of differentiation compared to control cultures. In the early phase of differentiation, this increase stemmed primarily from a rise in VE formation (Fig. 4B), and percentages of arterial (Fig. 4C) but not venous (Fig. 4C) VE increased in the presence of CHIR/SB. In addition, HOXA5, HOXA9 and HOXA10 transcription factors, previously shown to facilitate engraftment of iPSC-derived HSPCs^6^, were 2 to 4-fold more expressed in day 7 VE derived from CHIR/SB cultures compared to controls. However, the CHIR/SB-mediated effect on the production of VE was not sustained after day 7 of culture (Fig. 4B), and CD144^-^CD34^-^ mesenchymal cells continued to account for up to 95% of the adherent monolayer in the late phase of differentiation (Fig. 4F).

**Figure 4.**
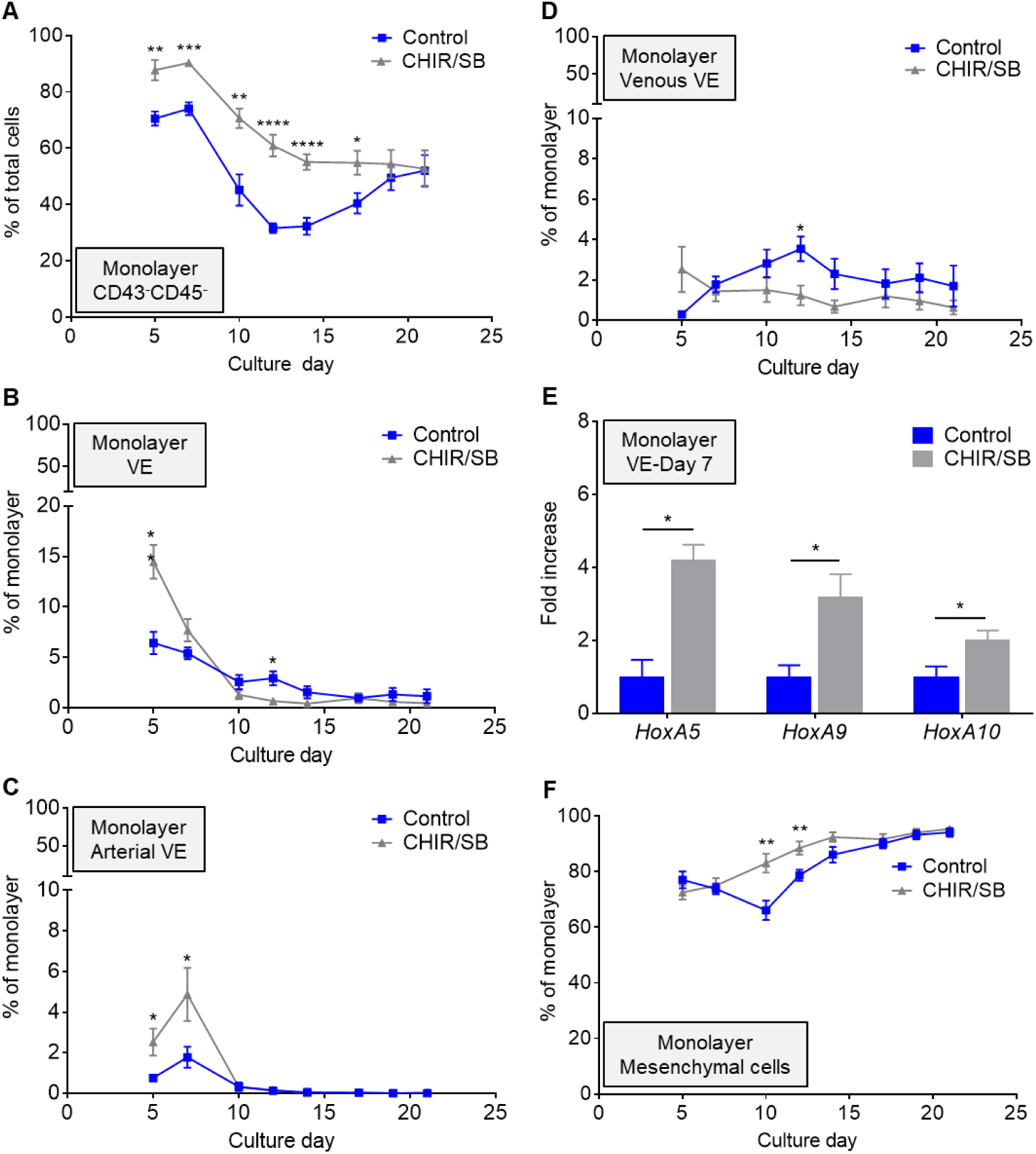
CHIR/SB molecules increase arterial VE in the adherent monolayer during the early phase of human iPSC differentiation. (**A**) Percentage of CD43^-^CD45^-^ monolayer cells in control and CHIR/SB-supplemented cultures at various days of differentiation (n=8). (**B**) Percentage of CD34^hi^CD144+ VE cells within the monolayer fraction of control and CHIR/SB-supplemented cultures at various days of differentiation (n=7). (**C**) Percentage of CD73^mid^CD184+ arterial VE cells within the monolayer fraction of control and CHIR/SB-supplemented cultures at various days of differentiation (n=7). (**D**) Percentage of CD73^hi^CD184^-^ venous VE cells within the monolayer fraction of control and CHIR/SB-supplemented cultures at various days of differentiation (n=5). (**E**) Fold increase in HOXA gene expression measured by real-time qPCR in CD34^hi^CD144+ VE cells sorted from CHIR/SB-supplemented cultures at D7 of differentiation relative to the same population sorted from control cultures (n=3). (**F**) Percentage of CD34^-^CD144^-^ mesenchymal cells within the monolayer fraction of control and CHIR/SB-supplemented cultures at various days of differentiation (n=5). Results are displayed as mean ± SEM; *P<0.05, **P < 0.01, ***P < 0.001, ****P<0.0001, by two-way unpaired Student’s t-tests comparing percentages of cells (A-D and F), or D7 untransformed ΔCT values (E) in control vs CHIR/SB groups at each culture day.

### 3.5 CHIR/SB molecules suppress primitive hematopoiesis and promote definitive HSPC formation during human iPSC differentiation

We next investigated whether the early increase in VE formation observed in the presence of CHIR/SB influenced hematopoietic development. Addition of CHIR/SB led to an overall decrease in the percentages (Fig. 5A) and numbers (Fig. S3B) of supernatant hematopoietic cells relative to control cultures. As reported in other systems^22^, we observed CHIR/SB-mediated suppression of the primitive wave of CD34^-^CD45^-^CD235a+ erythroid cells (Fig. 5B), and a rise in percentages of CD34^hi^CD45^lo^ definitive HSPCs (Fig. 5C) with a concomitant reduction in percentages of CD34^lo/-^ CD45+ cells in late stages of differentiation compared to controls (Fig. 5D). In colony forming assays, the frequency of progenitors with multilineage differentiation capacity was similar between sorted CHIR/SB and control supernatant cells at day 10 to 14 of culture (Fig. 5E). In contrast, the progressive rise in BFU-E/CFU-E frequency previously observed from day 17 to 21 of the standard differentiation protocol was further enhanced by addition of CHIR/SB (Fig. 5E). In RBCs differentiated from day 17 supernatant cells of CHIR/SB cultures, ε-globin expression was further decreased (5.8 ± 1.1% of total β-globin series expression) compared to controls (11.1 ± 2.1%), and a congruent increase in adult-type γ- and β-globins was observed (Fig. 5F).

**Figure 5.**
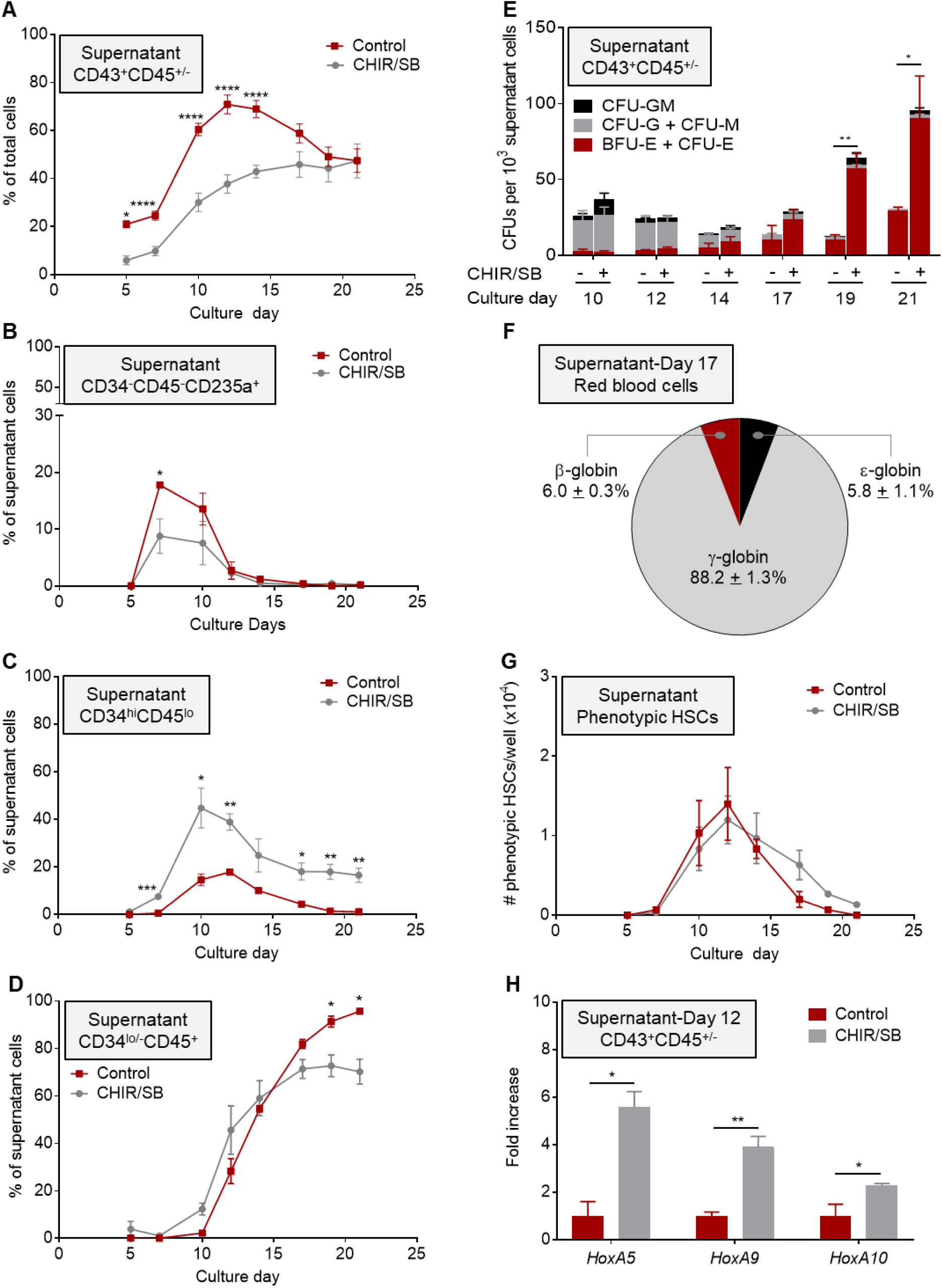
CHIR/SB molecules suppress primitive hematopoiesis and promote definitive HSPC formation during human iPSC differentiation. (**A**) Percentage of CD43+CD45+/^-^ supernatant cells in control and CHIR/SB-supplemented cultures at various days of differentiation (n=8). (**B**) Percentage of CD34^-^CD45^-^CD235a+ primitive erythroid cells within the supernatant fraction of control and CHIR/SB-supplemented cultures at various days of differentiation (n=5). (**C**) Percentage of CD34^hi^CD45^lo^ multilineage HSPCs within the supernatant fraction of control and CHIR/SB-supplemented cultures at various days of differentiation (n=5). (**D**) Percentage of CD34^lo/-^CD45+ hematopoietic progenitors within the supernatant fraction of control and CHIR/SB-supplemented cultures at various days of differentiation (n=5). (**E**) CFU counts per 1000 supernatant cells sorted from control and CHIR/SB-supplemented cultures at various days of differentiation (n=8). (**F**) Percentage of β-globin series expression in erythroid cells differentiated from D17 supernatant cells of CHIR/SB-supplemented cultures (n=2). (**G**) Absolute numbers of CD34+CD38^-^CD45RA^-^CD90+CD49f+ cells arising from 20-35 iPSC clusters (one well of a 12-well plate) in control and CHIR/SB-supplemented cultures at various days of differentiation (n=3). (**H**) Fold increase in HOXA gene expression measured by real-time qPCR in CD43+CD45^+/-^ supernatant cells sorted from CHIR/SB-supplemented cultures at D12 of differentiation relative to the same population sorted from control cultures (n=3). Results are displayed as mean ± SEM; *P<0.05, **P < 0.01, ***P < 0.001, ****P<0.0001, by two-way unpaired Student’s t-tests comparing percentages (A-D), CFU counts (E), cell numbers (G), or D7 untransformed ΔCT values (H) in control vs CHIR/SB groups at each culture day.

Next, we assessed whether CHIR/SB enabled the emergence of the third wave of definitive HSPCs with engraftment potential during iPSC differentiation. Phenotypically defined HSCs were readily detected at day 10 to 14 of differentiation at levels similar to control cultures (Fig. 5G). Addition of CHIR/SB also increased HOXA5, HOXA9 and HOXA10 expression 3 to 6-fold in supernatant cells harvested at day 12 compared to controls (Fig. 5H). However, these cells did not sustain long-term hematopoietic engraftment in NSG mouse recipients (Fig. S2B, right panel). Together, these results indicate that addition of CHIR/SB during mesodermal specification suppresses primitive hematopoiesis, promotes the second wave of definitive hematopoiesis, but is insufficient to enable generation of functional HSCs.

## 4. Discussion

Spontaneous production of cells with properties of normal transplantable HSCs, independent of ectopic expression of potentially oncogenic transcription factors, has to date eluded the field. In this study, we systematically characterized an alternative monolayer-based culture system that may facilitate investigation of novel strategies for the differentiation of human iPSCs into fully functional HSCs. The utility of this approach hinges on its scalability for clinical and research applications, and its simple, adaptable experimental design requiring no replating, EB formation or coculture on stromal elements.

In the supernatant fraction, we observed transient production of primitive CD235a+ erythroid cells followed by robust generation of HSPCs with clonogenic progenitor activity in vitro, a bona fide HSC immunophenotype (CD34+CD38^-^CD90+CD45RA^-^CD49f+), but no long-term engraftment potential in vivo. Thus, similar to other iPSC hematopoietic differentiation protocols, STEMdiff™ primarily recapitulates the first (primitive) and second (definitive) developmental wave of hematopoiesis observed in the yolk sac of developing embryos.

To understand the possible causes underpinning the absence of engraftable HSCs in this system, we examined the cellular constituents of the supportive adherent monolayer. We first identified a prevalent population of phenotypically defined mesenchymal cells throughout differentiation. Perivascular mesenchymal cells are known to interact with HSPCs and maintain their activity in the adult BM niche^42^. However, their role in promoting HSPC development during ontogeny has not been demonstrated. Instead, mesenchymal cells have been implicated as components of the niche in the yolk sac controlling primitive erythroid cell maturation^43^. Further investigation is needed to fully understand whether these cells may offer inhibitory signals that preclude normal developmental switch to third wave definitive hematopoiesis during iPSC differentiation. We also found limited VE production and arterial specification within the adherent monolayer that may account for the lack of engrafting HSCs in culture. Indeed, recent advances propose that formation of these cells is restricted to arterial vessels during ontogeny^16,17^. The arterial-specification model is supported in part by the demonstration of shared signaling requirements for both arterial identity acquisition and HSPC development^44-50^, and by the observation of selective impairment of hematopoiesis in the arteries of mice with knock-out of the artery-specific gene Ephrin B2^51,52^. Thus, in keeping with this model, arterial VE in the monolayer is likely inadequate to support the generation of bona fide engrafting HSCs in culture.

Our study provides proof-of-principle that the monolayer iPSC differentiation system is readily amenable to simple, clinically applicable modifications to improve hematopoietic output. Quantitative modulation of WNT/β-catenin and activin/nodal/TGFβ signaling pathways by one-time addition of CHIR/SB molecules during mesodermal specification enhanced arterial VE and increased HOXA gene expression and definitive HSPC formation. However, consistent with previous studies^20-22^, this approach alone was insufficient to orchestrate the formation of functional HSCs and additional revisions to this system will be required. Addition of retinoid acid, a morphogen with a documented role in de-repressing HOXA genes^53-56^ and in embryonic HSPC development^57,58^, may be required to complete the mesodermal patterning initiated by CHIR/SB. A recent study also revealed that overexpression ETS1, an EST family transcription factor involved in arterial fate control, or direct manipulation of MAPK/ERK signaling at the mesodermal stage led to an increase in arterial VE and definitive HSPCs^23^. It will be of interest to determine whether a combined approach to modulate MAPK/ERK, WNT/β-catenin and activin/nodal/TGFβ signaling pathways at this stage of differentiation can provide a synergy sufficient to further promote definitive hematopoiesis and the generation of engrafting HSCs.

In addition to the sequential emergence of developmental hematopoietic waves recognized in this study, our data also uncovered a nearly exclusive shift to definitive erythroid growth in later stages of differentiation that could be further enhanced by simple addition of CHIR/SB during mesodermal specification. Notably, this late erythropoietic upsurge occurred concomitantly with a rise in MSCs within the adherent monolayer. This observation coheres with the previously described erythroid commitment of human HSPCs co-cultured in the presence of MSCs^59,60^. In one study^60^, a significant enrichment of genes implicated in heme metabolism and a parallel downregulation of pathways involved in lymphoid and myeloid differentiation were noted in co-cultured CD34+ HSPCs. However, the precise molecular mechanism by which MSCs promote erythroid differentiation remain uncertain. The monolayer hematopoietic differentiation approach used in this study could provide a useful platform to identify prospective MSC-derived soluble factors implicated in erythroid maturation. Importantly, the STEMdiff™ system could be exploited to facilitate erythroid differentiation of iPSCs established from patients with hemoglobinopathies or various congenital bone marrow failures and anemias affecting early erythropoiesis. The robust generation of patient-specific erythroid progenitors could have considerable implications, including ex vivo modeling of disease pathophysiology, preclinical screening of gene therapy strategies, and specific testing of novel therapeutics against disease-relevant human cells.

## Acknowledgments

The authors thank J. Philip McCoy, Ph.D. and the NHLBI Flow Cytometry Core staff for sorting iPSC-derived cells; David Stroncek, M.D. and the NIH Department of Transfusion Medicine (DTM) and Cell Processing Section (CPS) staff for apheresis, selection and cryopreservation of human CD34+ cells; Tatyana Worthy, R.N., and the outpatient clinic nursing staff for recruiting normal volunteers and providing G-CSF administration teaching to healthy subjects; the mouse core facility staff for excellent animal care, Zu-xi Yu, M.D. Ph.D. of the pathology core of the NHLBI, NIH for sectioning and staining the teratomas. This work was supported by the intramural research program of the NHLBI, NIH. JPR was funded by a Howard Hughes Medical Institute (HHMI) Gilliam Fellowship, and this work was part of his Ph.D. thesis.

## Author contributions

JPR and AL conceived and designed the study. GC, MB and AL were involved in the development of the monolayer iPSC differentiation platform, patent filing and licensing to STEMCELL Technologies, Inc. JPR performed the majority of experimental procedures with help from HQ, HB and AL. JJHM performed erythroid differentiation experiments and globin expression analysis with general overview and helpful suggestions from NU and JFT. KK sorted the various cell populations. JZ and JB reprogrammed human cells to MCND-TENS2 iPSC lines, and confirmed pluripotency by flow cytometry. CL performed the teratoma assays. JPR and AL wrote and edited the manuscript, with contributions from all other authors.

