## Supplementary Figures and Tables for "Robust generation of erythroid and multilineage hematopoietic progenitors from human iPSCs using a scalable monolayer culture system"

(2) Translational Vascular Medicine Branch, NHLBI, NIH, Bethesda, MD 20892, USA.

(3) Clinical Flow Core Facility, NHLBI, NIH, Bethesda, MD 20892, USA.

(4) Transgenic Core Facility, NHLBI, NIH, Bethesda, MD 20892, USA.

(5) iPSC Core Facility, NHLBI, NIH, Bethesda, MD 20892, USA.

(6) College of Applied Medical Sciences, University of Hail, Hail, Saudi Arabia.

(7) Department of Biology, The Catholic University of America, Washington, DC 20064, USA.

**Author list footnotes:**

*** Corresponding author:** Andre Larochelle, M.D. Ph.D., National Heart, Lung and Blood Institute, National Institutes of Health, Bethesda, 9000 Rockville, Bethesda, MD 20892, USA.

☎ (301) 451-7139 (301) 496-8396

**This PDF file includes:**

Figures S1-S3
Tables S1 and S2

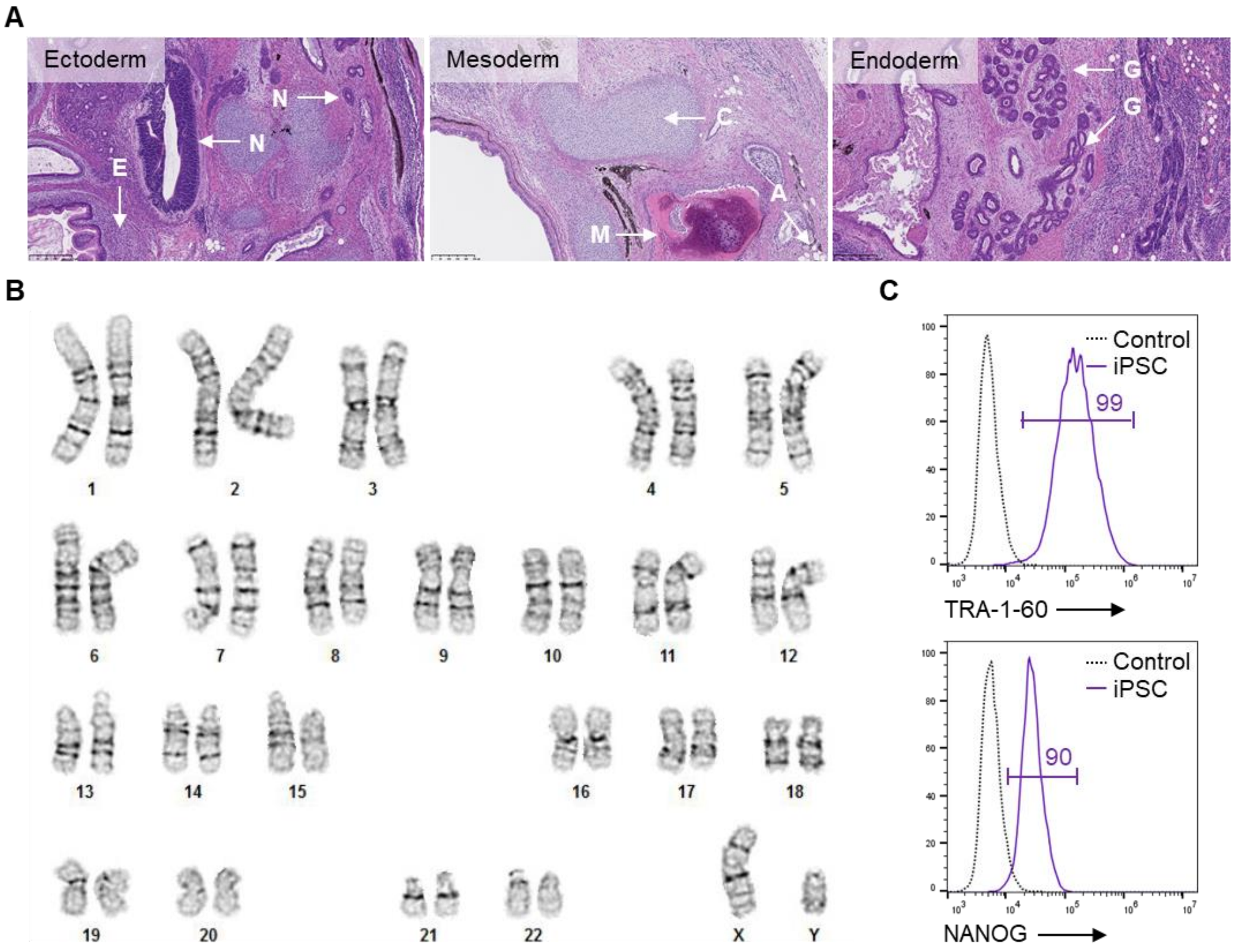


**Figure S1 | Characterization of MCND-TENS2 iPSC line.** (**A**) Hematoxylin and eosin stain of a teratoma from MCND-TENS2 iPSCs, displaying structures representative of ectoderm, mesoderm and endoderm. E, epidermal tissue; N, neutral tissue; M, striated muscle; C, cartilage; A, adipose tissue; G, gut epithelial tissue. Scale bar = 250 µm. (**B**) Cytogenetic analysis showing a normal karyotype (46, XY). (**C**) Flow cytometry analysis of iPSCs showing expression of pluripotency protein markers, TRA-1-60 and NANOG, compared to isotype controls.

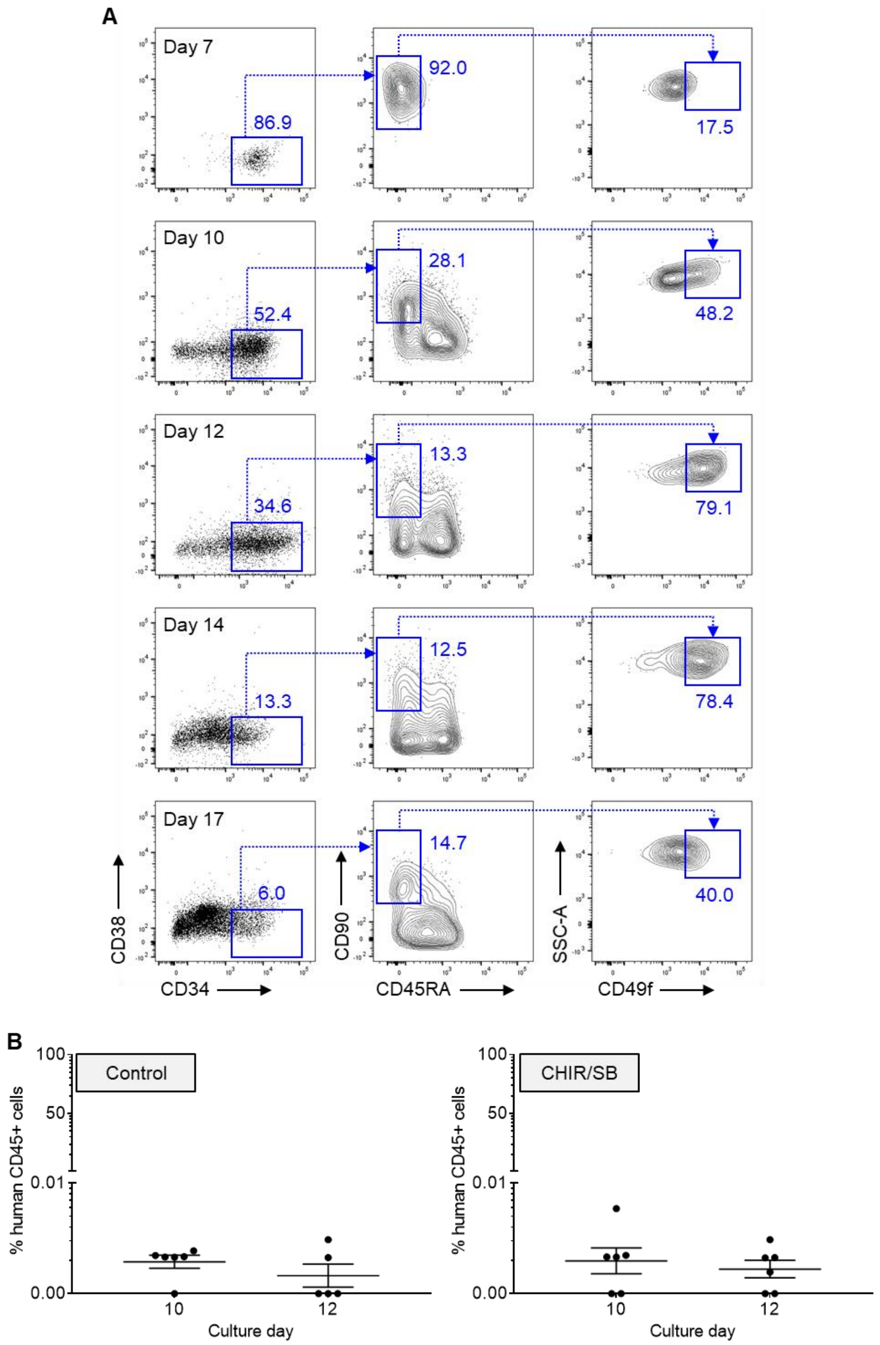


**Figure S2 | Production of phenotypically-defined HSCs with no long-term engraftment potential during hematopoietic differentiation of human iPSCs.** (**A**) Representative flow cytometry plots depicting $CD34^{+}CD38^{-}CD45RA^{-}CD90^{+}CD49f^{+}$ cells in gated $CD43^{+}CD45^{+/-}$ supernatant cells at various days of differentiation. (**B**) Human cell engraftment depicted as percentage of human CD45-expressing cells in the bone marrow of recipient NSG mice 16 weeks after transplantation of D10 or D12 sorted $CD43^{+}CD45^{+/-}$ supernatant cells derived from control (left panel, associated with Fig. 2E) or CHIR/SB-supplemented cultures (right panel, associated with Fig. 5G). Results are shown as mean ± SEM; each dot represents an individual mouse (n= 5 to 6 mice per group).

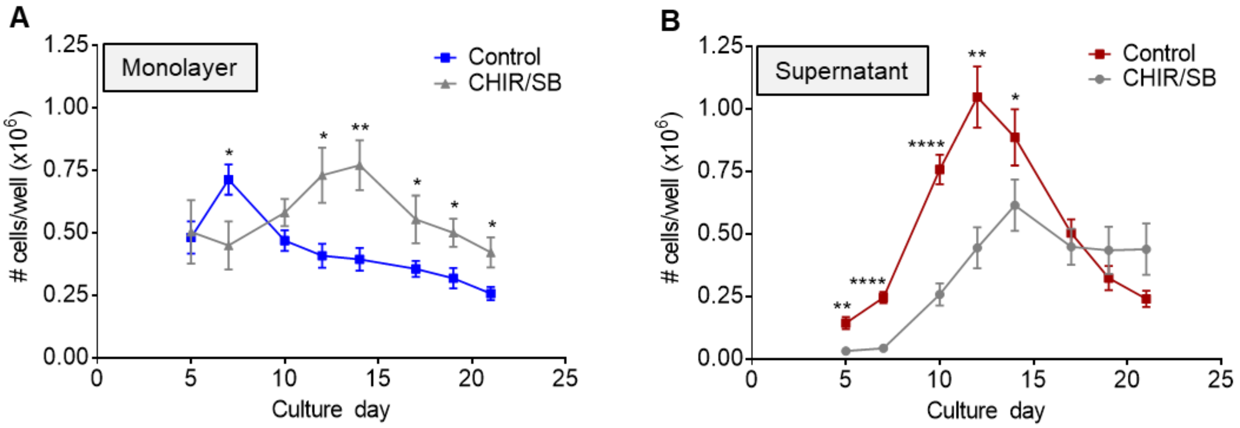


**Figure S3 | CHIR/SB molecules increase monolayer and decrease supernatant formation during hematopoietic differentiation of human iPSCs.** (**A**) Absolute numbers of $CD43^{+}CD45^{+/-}$ supernatant cells arising from 20-35 iPSC clusters (one well of a 12-well plate) in control and CHIR/SB-supplemented cultures at various days of differentiation (n=6). Associated with Fig. 4A. (**B**) Absolute numbers of $CD43^{-}CD45^{-}$ monolayer cells arising from 20-35 iPSC clusters (one well of a 12-well plate) in control and CHIR/SB-supplemented cultures at various days of differentiation (n=6). Associated with Fig. 5A. Results are displayed as mean ± SEM. *P<0.05, **P < 0.01, ****P<0.0001, by two-way unpaired Student's t-tests comparing cell numbers in control vs CHIR/SB groups at each culture day.

**Table S1 | Antibodies for flow cytometry analysis and FACS**

| **Antigen** | **Fluorochrome** | **Company/Catalogue #** | **Species** | **µL/test** |
|---|---|---|---|---|
| **CD34** | PE-Cy7 | BD Pharmingen 560710 | Mouse anti-human | 10 |
| **CD34** | PE | BD Pharmingen 555822 | Mouse anti-human | 20 |
| **CD38** | APC | BD Pharmingen 555462 | Mouse anti-human | 20 |
| **CD43** | BV711 | BD OptiBuild™ 743614 | Mouse anti-human | 5 |
| **CD43** | FITC | BD Pharmingen 555475 | Mouse anti-human | 20 |
| **CD45** | V450 | BD Horizon 560367 | Mouse anti-human | 5 |
| **CD45RA** | APC-H7 | BD Pharmingen 560674 | Mouse anti-human | 5 |
| **CD49f** | PE-Cy5 | BD Pharmingen 551129 | Rat anti-human | 20 |
| **CD73** | PE | BD Pharmingen 550257 | Mouse anti-human | 20 |
| **CD73** | FITC | BD Pharmingen 561254 | Mouse anti-human | 5 |
| **CD90** | PE-Cy7 | BD Pharmingen 561558 | Mouse anti-human | 5 |
| **CD105** | AF647 | BD Pharmingen 561439 | Mouse anti-human | 5 |
| **CD144** | BV605 | BD OptiBuild™ 743705 | Mouse anti-human | 5 |
| **CD184** | PE-CF594 | BD Horizon 562389 | Mouse anti-human | 5 |
| **CD235a** | FITC | BioLegend 349104 | Mouse anti-human | 5 |
| **N/A** | 7-AAD | Thermofisher 00-6993-50 | N/A | 5 |

**Table S2 | Taqman™ primers and probes**

| Gene | Assay or sequence | Probe |
|---|---|---|
| **GAPDH** | Life Technologies, Hs03929097_g1 | VIC |
| **HoxA5** | Life Technologies, Hs00430330_m1 | FAM |
| **HoxA9** | Life Technologies, Hs00365956_m1 | FAM |
| **HoxA10** | Life Technologies, Hs00172012_m1 | FAM |
| **ε-globin** | F: 5'-ACA ACC TCA AGC CCG C-3' | - |
| | R: 5'-AGA CAC CAG CTT CTG CC-3' | - |
| | P: 5'-HEX-TGC CAA AGT GAG TAG CCA GAA TAA TC-ZEN-IBFQ-3' | HEX |
| **γ-globin** | F: 5'-ACC TGG ATG ATC TCA AGG G-3' | - |
| | R: 5'-CAG TCA CCA TCT TCT GCC-3' | - |
| | P: 5'-Cy5-TGC CGA AAT GGA TTG CCA AAA CGG TC-IBRQ-3' | Cy5 |
| **β-globin** | F: 5'- ACA ACC TCA AGG GCA CC-3' | - |
| | R: 5'-ACA CCA GCC ACC ACT TTC-3' | - |
| | P: 5'-FAM- TGC CAA AGT GAT GGG CCA GCA CAC AG-IBRQ-3' | FAM |
| **α-globin** | F: 5'-TCC CCA CCA CCA AGA CCT AC-3' | - |
| | R: 5'-CCT TAA CCT GGG CAG AGC C-3' | - |
| | P: 5'-HEX-TCC CGC ACT TCG ACC TGA GCC A-IBRQ-3 | HEX |

F: forward primer; R: reverse primer; P: probe.
